# RBM3 enhances cold resistance by regulating thermogenic gene expressions

**DOI:** 10.1101/2023.05.10.540283

**Authors:** Junnosuke Nakamura, Megumi Uno, Takumi Taketomi, Manoj Kumar Yadav, Fuminori Tsuruta

## Abstract

Mammals are thermostatic animals capable of regulating their body temperature within a precise range, irrespective of ambient temperature conditions. However, the precise mechanisms by which a body temperature is controlled dependent on ambient temperature are still unclear. Here, we report that RNA binding motif protein 3 (RBM3), one of the cold-responsive proteins, regulates body temperature via expressing thermogenic genes during the late-postnatal period. The body temperature in *Rbm3* knockout (KO) juvenile mice was unstable and increased vulnerability to cold exposure. In addition, *Rbm3* KO mice exhibited increased lipid droplets in brown adipose tissue (BAT) and abnormal histology. The single-cell RNA-seq (scRNA-seq) analysis revealed that RBM3 is highly expressed in proliferating and differentiating cells in BAT. Moreover, RBM3 was necessary for upregulating thermogenic genes after cold shock. Notably, RBM3 interacted with UCP1 mRNA *in vivo*, thereby stabilizing its mRNA levels. Lastly, RBM3 regulated neuronal activity in the dorsomedial hypothalamic nucleus (DMH) under a cold environment. These data suggest that RBM3 regulates thermogenesis in juvenile mice through both upregulating thermogenic genes in BAT and activating DMH neurons after cold exposure.

## Introduction

Mammals possess physiological mechanisms to maintain a consistent body temperature range^1,2^. These mechanisms involve both heat production and dissipation, allowing adaptation to external temperature fluctuations. Maintaining an optimal body temperature is crucial for proper physiological functioning, including metabolic reactions. Conversely, deviations from the appropriate temperature range, such as fever and hypothermia, cause an abnormal physiological state, leading to multiple organ failures. Therefore, regulating body temperature is critical for maintaining a normal state.

When mammals are exposed to cold stimulation, heat production occurs in skeletal muscle and brown adipose tissue (BAT)^3,4^. Thermogenesis in skeletal muscle is particularly prominent in adult animals with well-developed musculature. In contrast, thermogenesis in BAT is pivotal for regulating body temperature during the postnatal period. Adipocytes are broadly classified into white adipocytes, which store excess energy as triacylglycerides in a single enlarged lipid droplet, and brown adipocytes, which contain multiple small lipid droplets and numerous mitochondria^5^. These mitochondria have uncoupling protein 1 (UCP1), which is involved in thermogenesis utilizing the energy of the proton gradient^6^. Thermogenesis due to changes in ambient temperature is dependent on the proliferation of preadipocytes and their differentiation into brown adipocytes, as well as increased expression of UCP1 in brown adipocytes. Previous studies have shown that cold stimulation differentiates from preadipocytes to brown adipocytes, increasing in these cells^7,8^. Ablation of sympathetic neurons projecting to BAT, which releases noradrenaline (NA), hampers the amplification of brown adipocytes in response to cold stimulation^9^, indicating that cold-sensitive neurotransmission from the center to the periphery is significant for adipocyte amplification and thermogenesis. Intriguingly, white adipocytes can transdifferentiate into beige adipocytes following cold shock^10^. Also, using an inducible labeling technique of mature adipocytes, beige cells have been reported to be differentiated from adipogenic precursor cells^11^. Because beige adipocytes have thermogenic functions like brown adipocytes, these differentiations are thought to improve the efficiency of heat production. The intriguing question is the molecular mechanisms underlying thermogenesis in the BAT. It is known that UCP1 expression is induced by the β-adrenergic receptor (βAR) via NA released from the sympathetic nerve. Shutting off adrenergic signaling results in a loss of resistance to cold stimulation^12–14^. As a downstream target of βAR, the protein kinase A (PKA) phosphorylates the cAMP response element binding protein (CREB). Previous study has reported that CREB binds to the promoter region in peroxisome proliferator-activated receptor γ co-activator-1α (PGC-1α)^15^. PGC-1α forms complexes with peroxisome proliferator-activated receptor γ (PPARγ) and these complexes play a significant role in expressing the *Ucp1* gene^16^. In addition, PKA has been known to activate p38 mitogen-activated protein kinase (p38 MAPK), leading to PGC-1α phosphorylation and subsequent UCP1 expression^17,18^. Furthermore, UCP1 expression associated with thermogenesis is triggered by activation of thyroid hormone receptors with triiodothyronine^19^. Therefore, UCP1 expression in BAT is mediated by multiple pathways, although some regulatory mechanisms remain to be elucidated.

So, how are the neural circuits involved in cold-induced heat production regulated? Previous studies have reported that the hypothalamus, which is a part of the central nervous system (CNS), plays a crucial role in controlling body temperature^20,21^. The preoptic area (POA) senses a change in ambient temperature, which is then integrated and transmitted the information to various effectors ^22^. Recently, inhibition of GABAergic neurons in the POA has been reported to activate the hypothalamic dorsomedial nucleus (DMH)^23^. The DMH activates a part of the brain called the rostral raphe pallidus (rRPa), which controls sympathetic neurons projecting to the BAT^24^. Apart from this neural circuit, it has been considered that the arcuate nucleus (ARC) and the ventromedial hypothalamus (VMH) in the hypothalamus also play a role in controlling the generation of heat in BAT^25,26^. Overall, it is likely that the thermogenic pathway involves a main route through POA-DMH-rRPa, as well as an additional pathway via the ARC, which influences not only heat production but also feeding behavior.

Recently, it has been revealed that exposure to cold stress can trigger the expression of cold-responsive proteins, including RNA binding motif protein 3 (RBM3). RBM3 is widely expressed in various tissues and is believed to modulate target mRNA translation by directly binding^27^. In addition, it has been shown that cold stress-induced expression of RBM3 promotes neurogenesis and proliferation of neural stem cells^28–30^. Importantly, the loss of the *Rbm3* gene causes abnormal cortical formation, indicating that RBM3 is crucial in modulating brain development^29^. Moreover, RBM3 has been reported to prevent neuronal cell death under neurodegenerative conditions^31^. RBM3 expression induced by cold stimuli attenuates degenerative phenotype in model mice such as Alzheimer’s disease (AD) and Creutzfeldt-Jakob disease (CJD)^31,32^. Thus, RBM3 is pivotal for regulating brain development and maintenance. Intriguingly, RBM3 is evolutionarily conserved only in mammals with a few exceptions^33^. RBM3 is also increased during hibernation^34,35^. Given that the body temperature significantly decreases during hibernation, RBM3 likely plays a role in controlling homeostasis, which in turn regulates body temperature. Although the relationship between cold shock and RBM3 expression has been studied, the detailed mechanism by which RBM3 is involved in controlling body temperature has not been clarified.

In this study, we report that RBM3 is a novel factor that regulates body temperature. We found that RBM3 is important in maintaining body temperature within the precise range. Also, RBM3 is required to suppress a decrease in body temperature after cold exposure. Importantly, RBM3 regulates the expression of thermogenic genes, including UCP1. Finally, RBM3 controls the neuronal activity in the DMH in response to cold exposure. These data suggest that RBM3 serves as a key factor that underlies regulating body temperature in a cold environment.

## Results

### Generation of Rbm3 knockout mice

To investigate the mechanisms by which RBM3 regulates body temperature *in vivo*, we generated *Rbm3* knockout (KO) mice using the improved-genome editing via oviductal nucleic acids delivery (*i*-GONAD) technique^36^. The *Rbm3* CRISPR RNAs (crRNAs) targeting the protospacer adjacent motif (PAM) in the first intron and fifth intron were delivered with recombinant Cas9 protein (1.0 μg/ml) and trans-activating crRNA (tracr RNA) into the embryo at day 0.7 of gestation in the oviductal ampulla by electroporation. Next, to confirm the genomic sequence of the *Rbm3*, we performed DNA sequencing and polymerase chain reaction (PCR). One mouse strain lacked 1.8 kbp of the genomic region spanning from the first to the fifth intron (Fig. 1A and 1B). We also verified the RBM3 expression using western blotting analysis. RBM3 was expressed in various tissues in WT mice and disappeared in *Rbm3* KO mice (Fig. 1C). The *Rbm3* KO mice were born at predicted mendelian ratios (Fig. 1D). Moreover, *Rbm3* KO mice exhibited more variability in body size and weight than WT mice at late-postnatal stage (P21), although their mean weight remained similar. On the other hand, there were no significant differences detected in average body size or weight at the adult stage (P60) (Fig. 1E and 1F).

**Figure 1.**
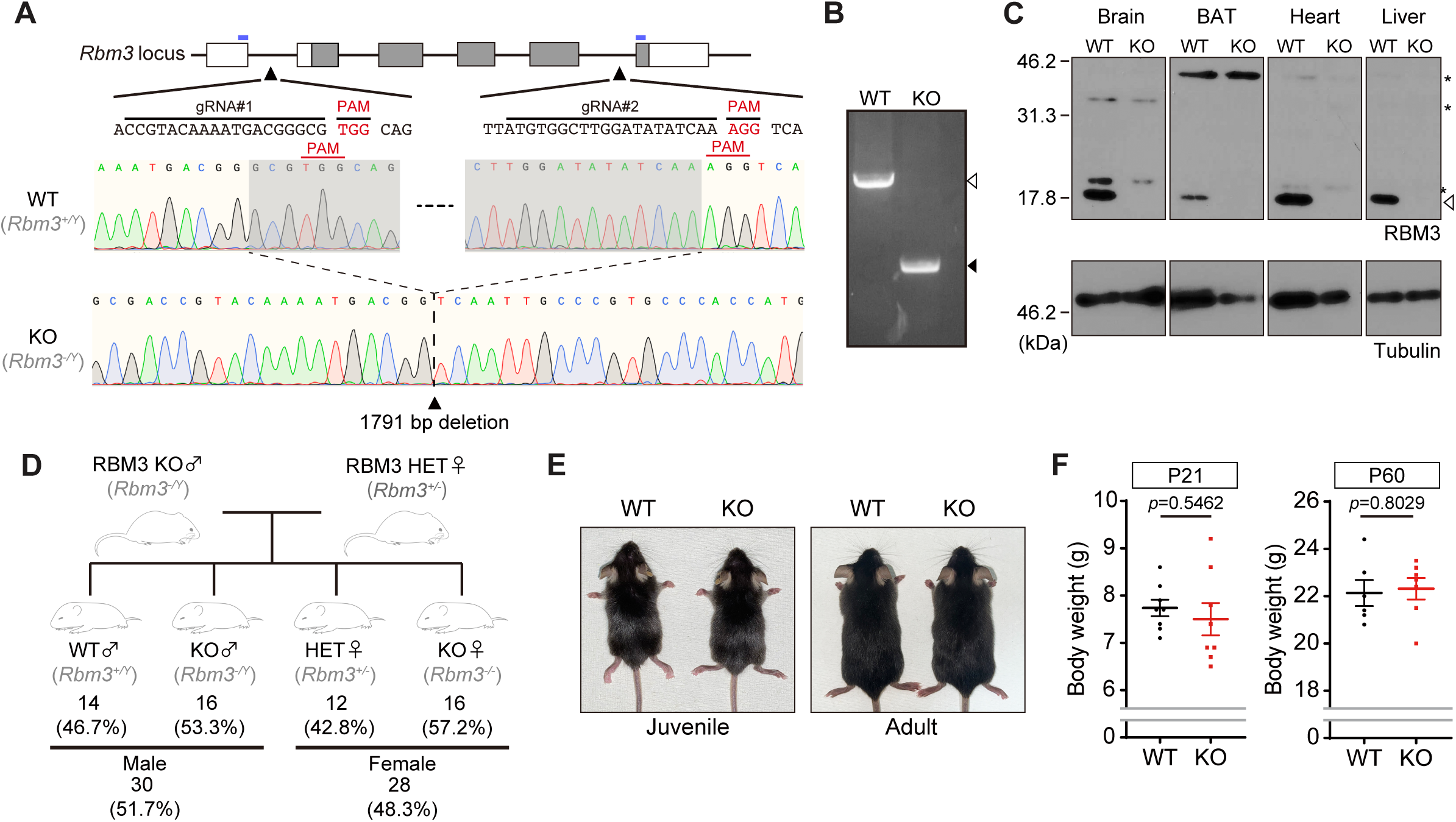
**(A)** Schematic diagram of the targeting strategy to eliminate the *Rbm3* gene. The guide RNAs (gRNAs) were designed to target the region between the first and second exons and between the sixth and seventh exons of the *Rbm3* gene (top panel). The waveform data for the sequencing results of PCR products amplified from F0 mice (bottom panel). RBM3^+/Y^: WT, RBM3^-/Y^: KO, The blue lines indicate primer recognition sites. **(B)** Genotyping analysis was performed on F0 mice with the expected fragment sizes of 2313 bp for WT (white arrowhead) and 522 bp for KO (black arrowhead). **(C)** Immunoblotting of RBM3 and Tubulin in the supernatant of indicated organs derived from WT and KO littermate mice at P21. The white arrowhead indicates RBM3 bands and the asterisks indicate the non-specific bands. **(D)** Genotype frequencies of postnatal mice produced from *Rbm3^-/Y^* and *Rbm3^+/-^* mice intercrosses. **(E)** Representative image of WT and *Rbm3* KO mice. Juvenile (P21) and adult (P60) littermates are shown. **(F)** The body weights of WT and *Rbm3* KO mice were measured at P21 and P60. P21 WT; n=8 mice, KO; n=8 mice, P60 WT; n=6 mice, KO; n=7 mice, mean ± SEM. *p* value was analyzed by Student’s *t*-test.

### RBM3 inhibits a decrease in body temperature after cold stimulation

To examine whether RBM3 is necessary for maintaining body temperature, we measured the rectal temperature in juvenile (P21) and adult (P60) mice. The average body temperature did not differ between the WT and *Rbm3* KO juvenile mice, whereas the variance in body temperatures was significantly higher in KO mice than in the WT mice [*p*=0.0089 (*F* test)]. On the other hand, there were no significant differences between WT and KO at the adult stage [*p*=0.9858 (*F* test)] (Fig. 2A), indicating that RBM3 is required for controlling body temperature within the appropriate range under the late-postnatal stage.

**Figure 2.**
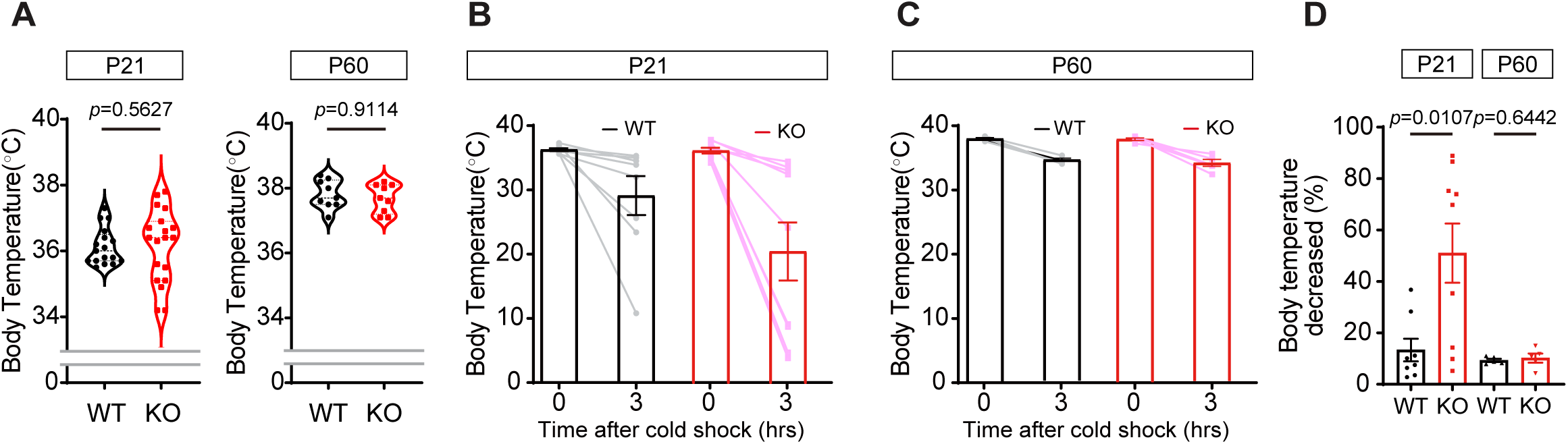
**(A)** The core body temperatures of WT and *Rbm3* KO mice were measured at P21 and P60 at room temperature (25°C). P21 WT; n=17 mice, KO; n=19 mice, P60 WT; n=9 mice, KO; n=9 mice, mean ± SEM. *p* value was analyzed by Student’s *t*-test. **(B)** The core body temperatures of WT and *Rbm3* KO mice (P21) were measured before and after a cold stimulation (4°C). WT; n=8 mice, KO; n=9 mice, mean ± SEM. *p* value was analyzed by Student’s *t*-test. **(C)** The core body temperatures of WT and *Rbm3* KO mice (P60) were measured before and after a cold stimulation (4°C). WT; n=5 mice, KO; n=5 mice, mean ± SEM. *p* value was analyzed by Student’s *t*-test. **(D)** Rate of decrease in body temperature in a cold environment. mean ± SEM. *p* value was analyzed by Student’s *t*-test.

Because RBM3 is a cold-induced protein, we analyzed whether RBM3 controls the body temperature in response to cold shock exposure. The body temperature in adult mice was not largely reduced by cold shock stimulation, regardless of loss of the *Rbm3* gene. However, the body temperature in juvenile mice was fragile against cold shock in *Rbm3* KO mice (Fig. 2B to 2D). These data suggest that RBM3 is a crucial protein that prevents a reduction of body temperature by cold shock exposure in juvenile mice but not in adult mice.

### RBM3 increases thermogenic gene expressions

Since BAT is known to be a heat-producing organ, we next investigated BAT abnormalities in *Rbm3* KO mice. First, we observed BAT stained with hematoxylin and eosin (HE). Similar to KO mice of other thermogenic genes^37^, BAT showed enlarged lipid vacuoles in *Rbm3* KO mice; however, the number of vacuoles was little different (Fig. 3A to 3C). We next explored the cells in the BAT where RBM3 is expressed. BAT contains immune cells, such as macrophages and dendritic cells, and also non-immune cells, such as vascular endothelial cells and adipose stem cells in addition to brown adipocytes (Fig. 3D). In addition, recent studies have reported that macrophages in BAT have a pivotal role in thermogenesis by regulating UCP1 expression in brown adipocytes^38^. Since it was technically difficult to identify the cell population in which RBM3 is expressed by BAT immunostaining, we used the single-cell RNA-seq (scRNA-seq) database to analyze the RBM3-expressing cell population. Analyses of the stromal vascular fractions (SVF) with or without cold stimulation suggested that RBM3 is expressed in almost all cells, but it was particularly expressed in proliferating and differentiating cells in non-immune cells (Cluster 2 and Cluster 3) (Fig. 3E and 3F). Importantly, the total number of RBM3-expressing cells was slightly increased after cold stimulation in non-immune cells (Fig. 3G). As these proliferating and differentiating cells differentiate into brown adipocytes^8^, it is likely that RBM3 is expressed in these subcellular populations which contributes to heat production. In contrast, in immune cells, macrophages with low expression of RBM3 were increased after cold stimulation, implying that subpopulations of macrophage-expressing RBM3 were diversified (Cluster 6 and Cluster 7) (Fig. 3H and 3I). Moreover, the bulk RBM3 expression level was slightly reduced after cold stimulation (Fig. 3J). Based on these *in silico* analyses, we concluded that the RBM3-expressing cells in BAT involved in thermogenesis are mainly brown adipocytes. Whereas, we speculated that cold stimulation to macrophages contributes to the proliferation rather than promoting RBM3 expression.

**Figure 3.**
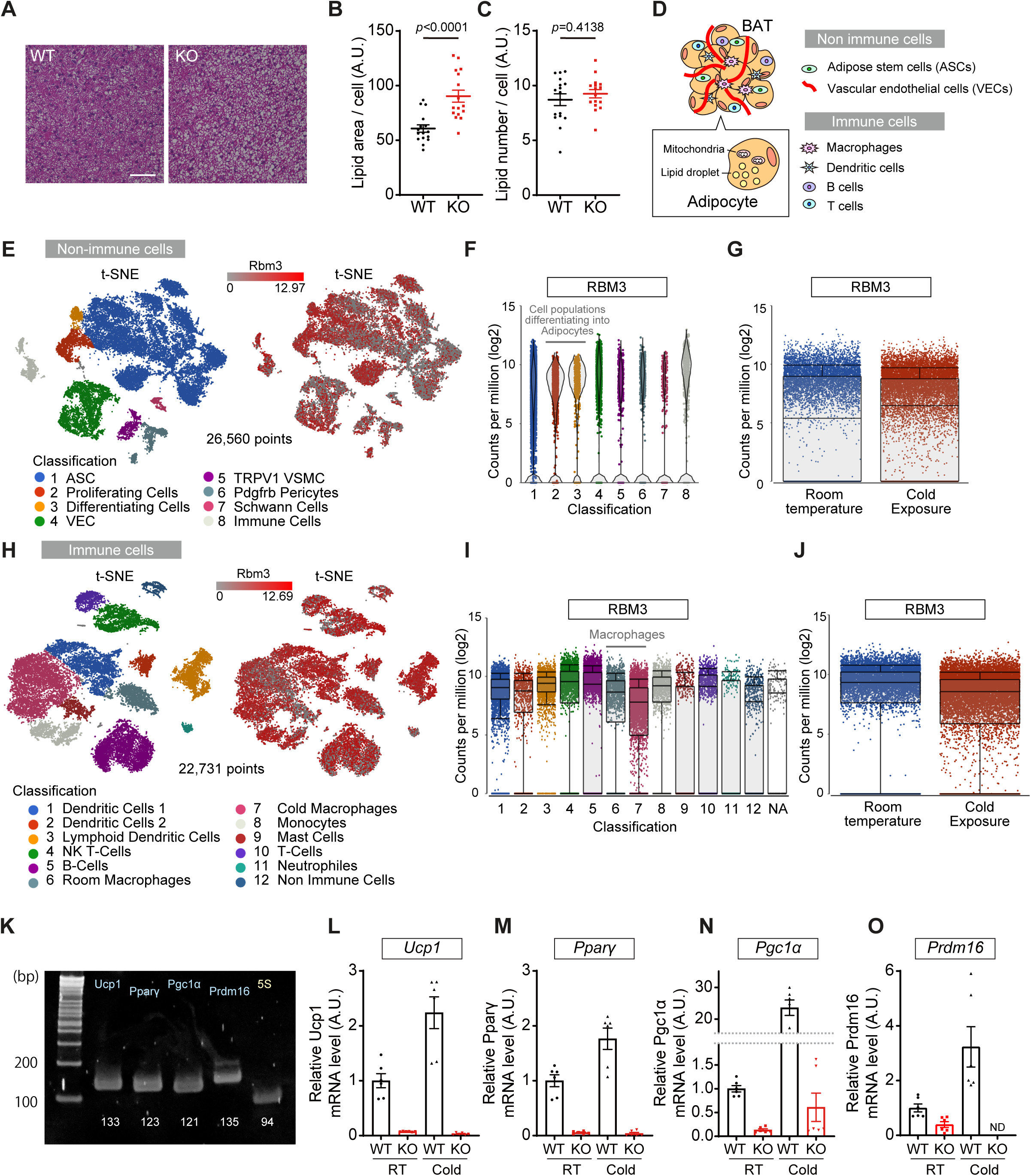
**(A)** Representative HE staining of the BAT obtained from WT and *Rbm3* KO mice (P21). Scale bar: 50 μm. **(B)** The size of the vacuole in BAT of WT and *Rbm3* KO mice per cell was measured using the particle analyzer by FIJI ImageJ software. 8 images were obtained from each mouse (n=16 images, 2 independent mice), mean ± SEM. *P* value was analyzed by Student’s *t*-test. **(C)** The lipid numbers in BAT of WT and *Rbm3* KO mice were measured using the particle analyzer by FIJI ImageJ software. 8 images were obtained from each mouse (n=16 images, 2 independent mice), mean ± SEM. *p* value was analyzed by Student’s *t*-test. **(D)** Schematic diagram showing the cells constituting brown adipose tissue. **(E)** The scRNAseq of non-immune cells from stromal vascular fraction of BAT. t-SNE plot showing the classification of the cells and Rbm3 expression in the cell population. **(F)** Violin plots of RBM3 expression levels of individual clusters from (E). **(G)** Rbm3 expression is increased after cold exposure. **(H)** The scRNAseq of immune cells from stromal vascular fractions of BAT. t-SNE plot showing the classification of the cells and Rbm3 expression in the cell population. **(I)** Boxplots of RBM3 expression levels of individual clusters from (H). **(J)** Rbm3 expression is decreased after cold exposure in immune cells. **(K)** Thermogenic gene (*Ucp1*, *Ppar*γ, *Pgc1*α, and *Prdm16*) mRNA and control (5S) expression of BAT was analyzed by RT-PCR. Each number indicates the size of PCR products (bp). **(L to O)** The mRNA of each thermogenic gene in the BAT of WT and *Rbm3* KO mice. The RNAs were isolated from the BAT (P21) in the presence or absence of cold stimulation for 3 hours. 5S rRNA levels were used for normalization. 2 samples were obtained from each 3 mice BAT (total n= 6), mean ± SEM. *p* value was analyzed by Student’s *t*-test. N.D., not detected.

We next examined whether RBM3 is necessary for maintaining thermogenic gene expression in response to cold shock. Firstly, we verified the primer for the thermogenic genes, such as *Ucp1*, *Ppar*γ, *Pgc1*α, and *Prdm16*, and confirmed whether these genes are expressed in the BAT of juvenile mice (Fig. 3K). All thermogenic genes are upregulated in response to cold stimulation; however, the expression level of these genes markedly reduced in *Rbm3* KO mice BAT (Fig. 3L to 3O). These data suggest that RBM3 is necessary for upregulating thermogenic genes in response to cold stimulation.

### RBM3 interacts with Ucp1 mRNA

To investigate whether RBM3 directly regulates these thermogenic genes, we searched the RBM3 recognition motif in the transcripts (Fig. 4A)^39^. The RNA binding domain of RBM3 resided in the N-terminal region (Fig. 4B). We examined the entire region of the mature mRNAs and found that all genes have RBM3 binding motifs (Fig. 4C and Suppl Table 1). Thereby, we examined whether RBM3 interacts with these transcripts using endogenous RNA immunoprecipitation (RIP) assay. We dissected mice, extracted BAT, and prepared samples for RIP analyses (Fig. 4D). Immunoprecipitated RBM3 was detected in WT but not in control IgG and KO samples (Fig. 4E). Using these precipitated samples, we conducted qPCR analyses. Consequently, *Ucp1* mRNA co-immunoprecipitated with endogenous RBM3 was significantly increased compared to controls. An increasing trend was also observed for *Ppar*γ mRNA. On the other hand, no significant differences were observed in *Pgc1*α and *Prdm16* mRNAs (Fig. 4F to 4I). These data suggest that RBM3 stabilizes *Ucp1* and possibly *Ppar*γ mRNAs directly and also upregulates other thermogenic genes through another machinery.

**Figure 4.**
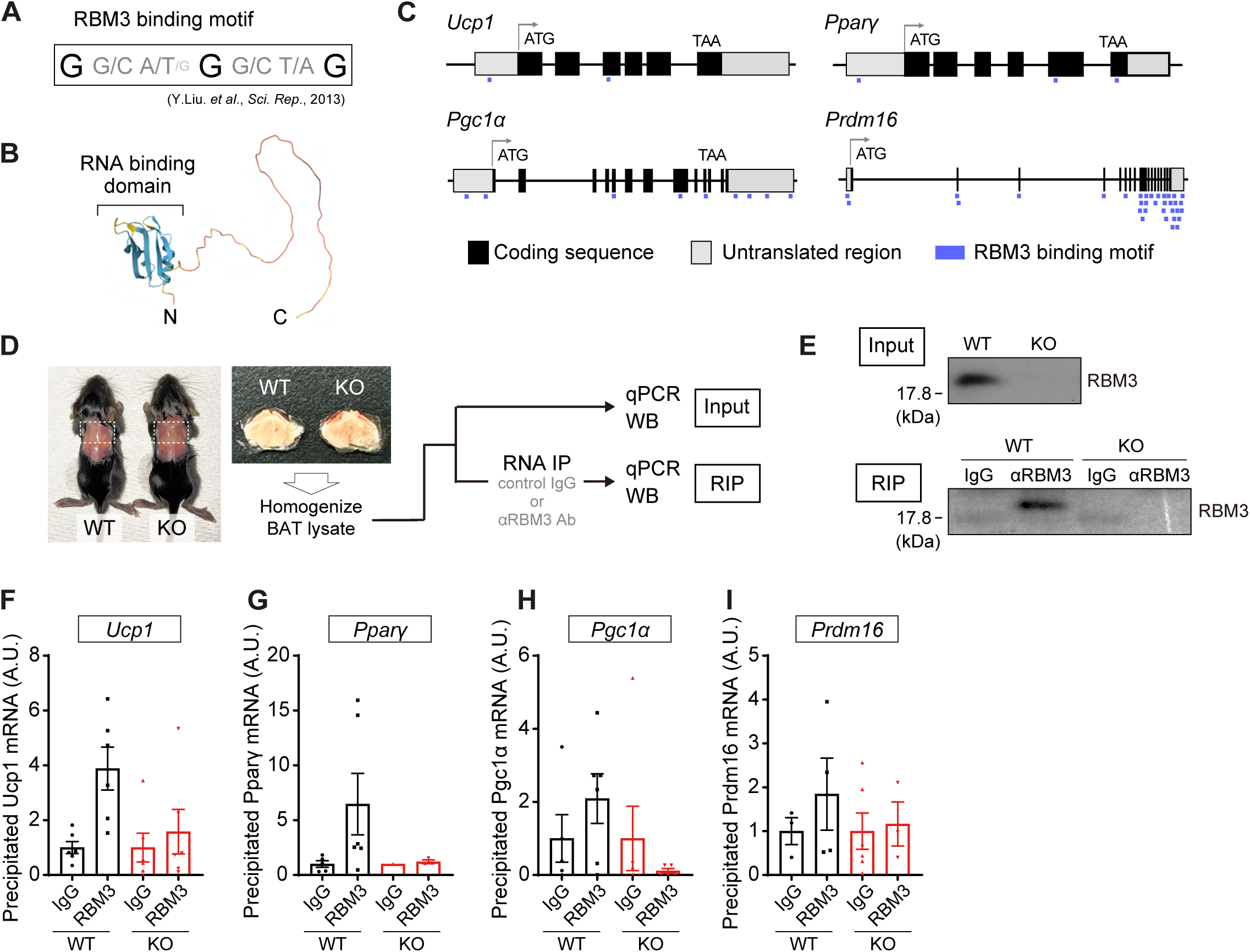
**(A)** Consensus sequence of RBM3 binding motif. **(B)** Schematic diagram showing the modeled structure of the RBM3 protein (AlphaFold: AF-O89086-F1) **(C)** Schematic diagram of exon-intron structure for thermogenic genes. The blue lines indicate the putative RBM3 binding sites. **(D)** The experimental flow of RNA immunoprecipitation (RIP) assay. **(E)** Validation of endogenous RBM3 immunoprecipitated with anti-RBM3 antibody. **(F to I)** TheqPCR analyses each thermogenic gene in precipitated samples. The input ΔCt values were used for normalization. 2 samples were obtained from each 3 mice (total n= 6), mean ± SEM. *p* value was analyzed by Student’s *t*-test.

### RBM3 is required for cold-induced neuronal activity in the DMH

Despite the markedly reduced expression of thermogenic genes in *Rbm3* KO BAT (Fig. 3L to 3O), *Pgc1*α and *Prdm16* mRNAs were not precipitated with RBM3 (Fig. 4H and 4I). These data prompt us to investigate whether RBM3 regulates not only working in BAT but also in the upstream, such as a thermosensory neural circuit. To test this idea, we focused on DMH and VMH in the hypothalamus, which are important for thermoregulation, as well as ARC, which is involved in feeding behavior. ARC is known to control not only feeding behavior but also thermogenesis^20,40,41^ (Fig. 5A). No abnormality in the hypothalamic region was observed in *Rbm3* KO mice (Fig. 5B). We then investigated whether RBM3 is necessary for neuronal activity under a cold environment. To examine this, we stained c-fos, which is one of the immediate early genes, in the hypothalamus. Importantly, c-fos expression was clearly increased in the DMH following cold shock; however, this effect was canceled in the KO mice brain. Whereas other regions, such as VMH and ARC, had little effect on the expression of c-fos (Fig. 5C to 5F). These data suggest that RBM3 in the brain is crucial for regulating neural circuits for thermogenesis in the hypothalamus.

**Figure 5.**
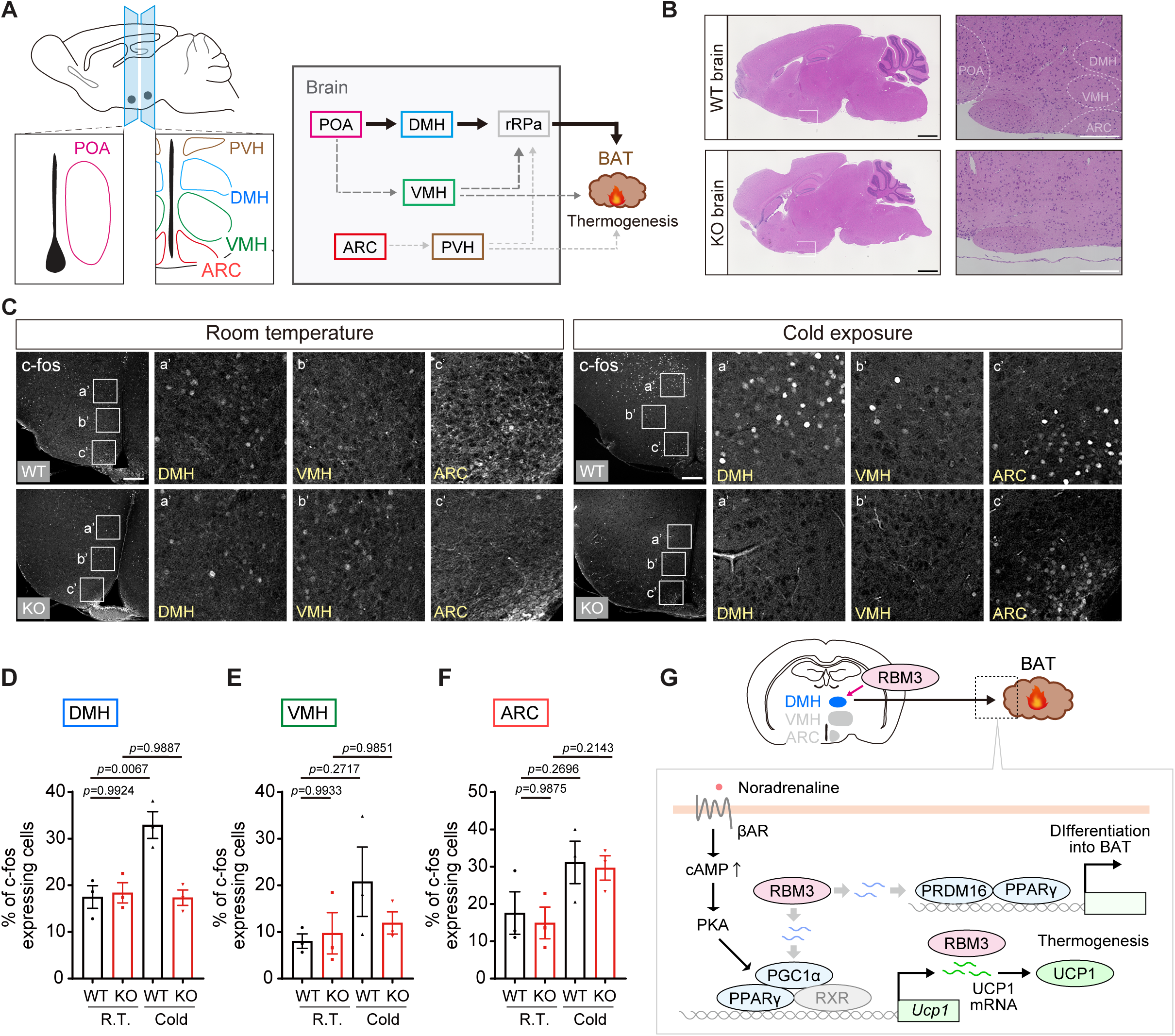
**(A)** Schematic diagram showing the neural circuit in the hypothalamus involved in the thermogenesis. **(B)** Representative HE staining of the brain was obtained from P21 WT and *Rbm3* KO mice. Left scale bar: 1000 μm, right scale bar: 200 μm. **(C)** Immunostaining in the hypothalamic region was obtained from WT and *Rbm3* KO mice (P21) in the presence or absence of cold stimulation (4°C, 3 hours). Scale bar: 200 μm. DMH: dorsomedial hypothalamic nucleus. VMH: ventromedial hypothalamic nucleus. ARC: arcuate nucleus. **(D∼F)** Quantification of c-fos expression level in the DMH, VMH, and ARC. Each sample was obtained from each 3 mice, mean ± SEM, *p* value was analyzed by one-way analysis of variance (ANOVA) Tukey–Kramer statical tests. **(G)** The model describing the function of RBM3 involved in cold-induced thermogenesis in the BAT.

## Discussions

Here, we investigated the role of RBM3 in thermogenesis under a cold environment. We found that RBM3 deficiency causes dispersion of body temperature in juvenile mice, rendering them vulnerable to cold stimuli. In addition, RBM3 was required for upregulating thermogenic genes in BAT and activation of DMH neurons under a cold environment. Overall, these findings provide a new mechanism for RBM3-dependent thermogenesis in juvenile mice (Fig. 5G).

The homeostatic mechanisms responsible for regulating body temperature in mammals are established with growth, which means regulating body temperature during postnatal stages is unstable. Furthermore, a significant deviation from body temperature will result in death, making the maintenance of body temperature during this period critical for sustaining life. However, the mechanisms required to maintain body temperature during this period were still unclear. Previously, it has been reported that RBM3 expression increases after birth in the rat cerebellum but decreases as development progresses, becoming constant at about 6 to 8 weeks after birth^42^. Moreover, we observed that juvenile mice lacking RBM3 show unstable body temperature, while adult mice show normal body temperature. Indeed, our preliminary data have suggested that skeletal muscle shivering is not crucial for cold shock-induced thermogenic genes in neonates (unpublished data). It seems that brown adipocytes develop during the neonatal period and subsequently shrink with growth in hamsters^43^. Therefore, it is likely that RBM3 contributes to maintaining body temperature during the neonatal period by regulating BAT heat production. Importantly, the pathological findings of lipid droplets in *Rbm3* KO BAT were similar to those in *Ucp1* KO BAT^37^. The proton gradient mediated by the β-oxidation and glycolysis contributes not to ATP synthesis but to heat production through UCP1 activity. Since UCP1 expression is reduced in *Rbm3* KO mice, fatty acids and glucose might accumulate in lipid droplets and enlarge due to reduced metabolic pathways of β-oxidation and glycolysis. Since expression of thermogenic genes, including *Ucp1*, is reduced in *Rbm3* KO mice, the metabolic pathways of β-oxidation and glycolysis may be reduced as negative feedback, resulting in fatty acid and glucose accumulation and enlarged lipid droplets.

Recent studies have shown that RBM3 regulates the stability and translation of the target mRNA via direct binding (G[CG][ATG]G[GC][TA]G)^39^. Thus, it is likely that RBM3 acts as the regulatory factor for thermogenesis via modulating target mRNAs in a context-dependent manner. Indeed, we found that thermogenic genes (*Ucp1*, *Ppar*γ, *Pgc1*α, and *Prdm16*) in BAT are clearly decreased in *Rbm3* KO mice.

Particularly, RBM3 directly interacted with *Ucp1* mRNA and possibly *Pparg* mRNA *in vivo*. Given that RBM3 regulates heat production in BAT, its function should be considered not only in adipocytes but also in other cell types. Previous studies have shown that loss of MAFB in macrophages increases IL6 secretion, leading to a proinflammatory response in BAT, reduced sympathetic innervation, and decreased UCP expression and heat production in adipocytes^38^. Because RBM3 is also expressed in macrophages (Fig. 3H and 3I), elucidating the biological relevance of RBM3 in MAFB is an interesting question. Furthermore, it has been reported that Calsyntenin 3β (CLSTN3β) in adipocytes stimulates S100β secretion and enhances NA release from sympathetic neurons^44^. S100β in the CNS is expressed in astrocytes and oligodendrocytes. These observations suggest that RBM3 might promote the *S100*β gene expression in Schwann cells, leading to UCP1 expression in adipocytes.

RBM3 has been demonstrated to directly bind to and regulate the expression of thermogenic genes, such as *Ucp1* mRNA, in BAT. In contrast, while RBM3 enhanced *Pgc1*α and *Prdm16* mRNA levels in BAT, RBM3 did not bind to these mRNAs directly. These observations suggest that RBM3 may regulate thermogenic machinery in not only BAT but also other tissues, such as the brain, through non-tissue-autonomous mechanisms. In this study, we revealed that RBM3 is necessary for DMH activation after cold shock. Hypothalamic control of body temperature is regulated by temperature-sensitive neurons in the POA. Under normal conditions, GABAergic neurons in the POA project inhibitory signals to the DMH; however, cold stimulation reduces GABAergic inhibition from the POA to the DMH. Subsequently, glutamatergic neurons in the DMH activate rRPa neurons, transmitting signals to BAT for heat production^45^. Indeed, activation of DMH neurons by optogenetics enhances heat production^24,46^. Thus, DMH functions as a crucial neuronal nucleus in response to low-temperature stimulation. Because loss of the *Rbm3* gene attenuates DMH neuronal activity in response to cold stimulation, the potential targets in the DMH neurons or the POA neurons may exist. For instance, the neuropeptide Y (NPY) in the DMH has been reported to control the expression of *Ucp1* mRNA in the BAT^47^. As NPY1r, its receptor protein, has five RBM3 binding motifs, it is possible that RBM3 in the CNS regulates the expression of NPY1r expression. Moreover, Nrd1, a metalloendopeptidase, is secreted and regulates extracellular conditions. A defect in the *Nrd1* gene decreases the UCP1 expression in response to cold stimulation and impairs the body temperature control^48^. As Nrd1 mRNA also has more than ten RBM3 binding motifs in the 5’UTR and 3’UTR, RBM3 may directly regulate Nrd1 expression in neurons, subsequently increasing the expression of UCP1 in the BAT.

It has been reported that RBM3 is involved not only in regulating body temperature but also in controlling circadian rhythms. Knockdown of the Rbm3 gene disrupts the expression of circadian genes such as Dbp and Per2. It is also proposed that RBM3 stabilizes these mRNAs through potentially direct binding^39^. Moreover, RBM3 regulates rhythmic neural activity during the day/night cycle in cultured hippocampal neurons^49^. Thus, RBM3 is tightly involved in circadian rhythms. A key question is the biological relevance between body temperature and circadian rhythm. Recent intriguing research has revealed that the rhythmic expression of RBM3 depends on BMAL and regulates neural circuits in the suprachiasmatic nucleus (SCN), the control center for circadian rhythm regulation^50^. This suggests that the BMAL-RBM3 axis plays a crucial role in integrating temperature and circadian rhythm control. In this study, we found that cold stimulation induces RBM3 expression in the DMH. Given that body temperature changes rhythmically, being higher during arousal and lower during sleep, the circadian rhythm-dependent changes in neural activity in the DMH present an intriguing and challenging area for future research.

In conclusion, our study has identified RBM3 as a novel factor that controls body temperature under normal conditions and demonstrated its influence on the expression of UCP family proteins. Additionally, we have shown that RBM3 is crucial for maintaining body temperature following cold stimulation. These findings shed light on the previously enigmatic machinery regulating body temperature and provide important insights into the connection between ambient temperature changes and the homeostatic state.

## Materials and Methods

### Mice

The C57BL/6N wild-type (WT) mice were obtained from Japan SLC, Inc. (Shizuoka Japan). The mice were maintained in standard conditions (Mice were housed in a 12 h: 12 h light: dark cycle with lights on at 8 am). All mice used in this research were male. All experiments were approved by the Animal Experiment Committee at the University of Tsukuba (the approval numbers: 19-340, 20-438, 21-443, 22-386, 23-370, 24-377) and conducted according to the university guidelines for animal care and use, and ARRIVE guideline. *Rbm3* KO mice of the C57BL/6N strain were generated by *i*-GONAD as previously reported^36^. Briefly, crRNA (Integrated DNA Technologies) was designed with the upstream-gRNA sequence: 5’-ACCGTACAAAATGACGGGCGTGG-3’, the downstream-gRNA sequence: 5’-ATGTGGCTTGGATATATCAAAGG-3’ to target the intron 1 and intron 5 of the *Rbm3* allele. In addition, 200 μM each Alt-R CRISPR Cas9 crRNA and Alt-R CRISPR Cas9 tracrRNA (Integrated DNA Technologies) were annealed at 94°C for 2 min, mixed with 1.0 ug/ml Alt-R S.p. Cas9 Nuclease (Integrated DNA Technologies), and injected into the oviduct at E0.7 of mice with confirmed plug according to the previous study. After injection, 40 V, 5 msec electrical pulses were given three times at 50 msec intervals, followed by 10 V, 50 msec electrical pulses three times at 50 msec intervals, using by gene delivery system NEPA21 (NEPA Gene). In order to fix the mutation in the strain, the F0 founder was mated with WT mice, and the resulting F1 heterozygous male and female mice were crossed to obtain an F2 generation containing homozygous mutants. The established strain was maintained by sibling breeding and used in the current study.

### Measurement of rectal temperature and body weight

The rectal thermometry of WT and *Rbm3* KO mice (P21 and P60) was measured using a digital thermometer (SATO KEIRYOKI, SK-1260). Briefly, the thermometer was inserted into the rectum, followed by a measurement of the temperature. The thermometry was conducted at 10 am on time. This rectal temperature is corresponding to the core body temperature^51^. This experiment was reproduced at least three times. For cold stimulation, mice were placed in cages without nestlets and left under normal conditions (room temperature) and cold environments (approximately 4∼8 °C) for 3 hours. The body weight of WT and *Rbm3* KO mice (P21 and P60) was measured using an electronic balance (METTLER TOLEDO, BD601). This experiment was conducted at 10 am on time.

### Immunoblot analysis

Samples were collected in homogenization buffer (20 mM Tris-HCl [pH 7.5], 150 mM NaCl, 1 mM EDTA pH8.0, 0.4% TritonX100, 1 mM DTT) and homogenized by DIGITAL HOMOGENIZER (As One International, Inc.). Lysates were centrifuged at 14,000 rpm for 5 min, and were then collected the supernatants. The concentration of samples was measured by the Bradford method (Coomassie Protein Assay Kit, Thermo Fisher). Adjusted samples were mixed with sample buffer (125 mM Tris-HCl pH 6.5, 4% SDS, 10% Glycerol, 0.01% bromophenol blue, 10% 2-mercaptoethanol) and were boiled at 100°C for 3 minutes. Equal amounts of protein (5 μg) were loaded into each well and analyzed by western blot analysis. After SDS-polyacrylamide gel electrophoresis, each sample was transferred to a polyvinylidene difluoride membrane (Pall Corp.). The membranes were blocked in 5% skim milk/TBST for 60 min at room temperature and were then incubated overnight at 4 °C with the primary antibodies [anti-RBM3 antibody (Proteintech, Cat#14363-1-AP, 1:1000) and anti-Tubulin antibody (SIGMA, DM1A, 1:1000)]. After washing with PBS, the membranes were incubated for 1 hour with the peroxidase-conjugated secondary antibodies (SeraCare, 1:20,000). The signals were detected by chemiluminescent substrate reagent (Chemi-Lumi One Super, Nacalai tesque)

### Quantitative RT-PCR

Brown adipose of WT or *Rbm3* KO mice were solved with 500 µl ISOGEN II (NIPPON GENE). The suspension was centrifuged at 12,000 x g for 15 min at 4°C. The colorless upper aqueous phase was collected, and added to 500 µl of 100% isopropanol, and centrifuged at 12,000 x g for 10 minutes at 4°C. The supernatant was removed, and the RNA pellet was washed in 75% ethanol and re-suspended in RNAase-free water. The reverse transcription was performed on 1 μg of total RNA using the solution containing 100 units ReverTra Ace (TOYOBO), 25[pmol Random Primer (TOYOBO), and 20[nmol dNTPs. The qPCR was performed in duplicate in a 96-well plate (Thermo Fisher Scientific) using THUNDERBIRD SYBR qPCR Mix (TOYOBO) in Applied Biosystems 7900HT Fast Real-Time PCR System (Applied Biosystems). Cycle threshold (CT) values were normalized to the 5S rRNA value. The relative quantity of the target expression was calculated by 2^-ΔΔCt^ methods using SDS Software 2.4.2 (Applied Biosystems) with the following calculation. The relative quantity[=[2^−ΔΔCt^, ΔΔCt = (Ct^target^ – Ct^5S^)_sample_ – (Ct^target^ – Ct^5S^)_reference_. See Supplementary Table 2 for statistical analysis of qRT-PCR.

### Immunohistochemistry

The brains collected from WT and *Rbm3* KO littermate mice were perfused with 4% PFA-PBS and fixed with the same fixative solution overnight. After 30% sucrose in PBS infiltration, the samples were embedded in Tissue-Tek OTC compound (SAKURA) and sliced at a 30 μm thickness by the cryostat (Leica Biosystems). Free-floating slices were permeabilized and blocked with PBS containing 0.25% Triton X-100 and 5% bovine serum albumin (BSA). Primary antibodies were incubated in PBS containing 0.1% Triton X-100 and 5% BSA for 1 day [anti-c-fos antibody (Abcam, Cat# ab208942, 1:1000)] at 4°C. Immunostaining was detected using Alexa 488 fluorescent secondary antibody (Thermo Fisher Scientific, 1:1000) with 5.0 μg/ml Hoechst 33342 (Invitrogen) incubated in PBS containing 5% BSA for 1 hour at room temperature. Slices were mounted in VECTASHIELD Mounting Medium (Vector Laboratories). After washing by PBS, the samples were observed using a confocal laser scanning microscope (LSM700, Carl Zeiss) with 10× (Plan-Apochromat 10×/0.8 M27) and 40× (Plan-Apochromat 40×/1.3 Oil DIC M27) objective. The diode excitation lasers (Diode405, and Diode555) were operated and directed to a photomultiplier tube (LSM T-PMT, Carl Zeiss) through a series of band pass filters (Ch1:BP420-475 + BP500-610, Ch2:BP490-635, and Ch3:BP585-1000).

### HE staining

The BAT collected from WT and *Rbm3* KO littermate mice were perfused with 4% paraformaldehyde in phosphate-buffered saline (4% PFA-PBS). The samples were fixed in the 4% PFA-PBS overnight, and embedded in paraffin. Paraffin sections were then sliced into 2 μm sections and stained with Meyer’s hematoxylin and eosin. Tissue specimens were observed using the fluorescence microscope (BZ-9000, Keyence) with 40x (PlanApo 40x/0.95) objective and a charge-coupled device camera (Keyence).

### *in silico* analysis of the scRNA-seq dataset

The scRNA-seq analysis was described previously^38^. Briefly, the scRNA-seq data of interscapular BAT was obtained from the public database (GSE207706)^8^ and analyzed using Partek Flow software (version 10.0.22.1111). Quality assessment and control measures were employed to exclude apoptotic cells, debris, empty reads, duplets, and batch effects. The data were then normalized and subjected to principal component analysis (PCA), graph-based clustering, and t-SNE plot visualization. Biomarkers were identified to characterize the cell populations, and Enrichr tools were used to define these populations based on various databases.

### RNA Immunoprecipitation

The BAT collected from WT and *Rbm3* KO littermate mice (P21) were homogenized in homogenize buffer (50 mM Tris-HCl [pH 7.5], 150 mM NaCl, 1 mM EDTA pH8.0, 0.5% NP40) containing complete mini protease inhibitor cocktail (Roche, Cat#04693124001) and RNase OUT (Invitrogen). Lysates were centrifuged at 14,000 rpm for 15 min and were then collected the supernatants. The supernatant was immunoprecipitated with either normal rabbit IgG (2 ng/μl) (Santa Cruz, sc-2027) or anti-RBM3 antibody (2 ng/μl) (Proteintech, Cat#14363-1-A) together with 50% slurry protein G-agarose beads (Thermo Fisher Scientific) at 4°C for 2 h. The beads were then washed with homogenize buffer and subjected to either immunoblot analysis or qRT-PCR after extraction of mRNA using Isogen II. All manipulations were performed as carefully as possible on ice to minimize mRNA degradation. The qRT-PCR was conducted in duplicate in a 96-well plate (Thermo Fisher Scientific) using THUNDERBIRD SYBR qPCR Mix (TOYOBO) in Applied Biosystems 7900HT Fast Real-Time PCR System (Applied Biosystems). Cycle threshold (CT) values were normalized to the input sample. The relative quantity of the target expression was calculated by 2^-ΔΔCt^ methods using SDS Software 2.4.2 (Applied Biosystems). Cycle threshold (CT) values were normalized to the input sample RNA value. The relative quantity of the target expression was calculated by 2^-ΔΔCt^ methods using SDS Software 2.4.2 (Applied Biosystems) with the following calculation. The relative quantity[=[2^−ΔΔCt^, ΔΔCt = (Ct^RIP^ – Ct^iuput^) – (Ct^RIP^ – Ct^input^) .

### Statistical analysis and image processing

The statistical analyses were calculated by using GraphPad Prism (GraphPad Software). All image processing was conducted by Adobe Creative Could CC 2024 (Photoshop 25.6.0 and Illustrator 28.5) and FIJI Image J 2.1.0/1.53c. To quantify the number and size of lipid droplets, the Analyze Particle function is Fiji/ImageJ software was used. The acquired HE stained slice images were converted to binarized images using the ’Li’ method to detect transparent areas. To identify the lipid droplets in contact with each other, the binarized images were processed using the ’Watershed’ function. To exclude blood vessels and detect only lipid droplets, the Analyze Particle function was set to detect particles with a size greater than 1 µm^2^, a circularity within the range of 0.6-1.0, and exclude those on the edges. To quantify the number of cells, HE stained images were split using the ‘Color Deconvolution’ function, and hematoxylin channel images were converted to binarized images using the ’RenyiEntropy’ method. the Analyze Particle function was set to detect particles with a size greater than 1 µm^2^, and exclude those on the edges. The c-fos expression in the DMH, VMH, and ARC were quantified using FIJI ImageJ. The area of nuclear staining was binarized for setting the region of interest (ROI). The number of c-fos and Hoechst33342 stained cells in the ROI were calculated using the Analyze Particle function. The size of ROI (10-150 μm^2^) was defined as a suitable area to extract the signals to eliminate the noise and meningeal cells. All images were taken with the same setting, including exposure time, signal amplification, and objective lens.

## Acknowledgments

We thank Drs. Motoshi Nagao (NRCD, Japan), Michito Hamada (Univ of Tsukuba), and the lab members for helpful discussions and technical supports. This work was supported by the Grant-in-Aid from the Ministry of Education, Science, Sports and Culture of Japan JSPS KAKENHI [20K05951 (FT)], Kao foundation for health science (FT), Gout and uric acid foundation (FT), JSPS Research Fellowship for Young Scientists [24KJ0482 (JN) and 23KJ0285 (TT)], JST SPRING [JPMJSP2124 (JN)], and partly supported by Center for Quantum and Information Life Sciences, University of Tsukuba (FT).

## Author contributions

J.N. and F.T. designed research; J.N., M.U. performed research; J.N., M.U., T.T., M.K.Y analyzed data; J.N and F.T. wrote the paper. Study supervision: F.T

## Notes

### Competing Interest Statement

The authors have declared no competing interest.

### Summary of Updates

Figure 1 and 2 updated; Figure 3, 4, and 5 added as new data; authors updated; Supplemental tables updated.

