## Supplemental Tables for "RBM3 enhances cold resistance by regulating thermogenic gene expressions"

**Supplementary Table 1: RBM3 binding site in thermogenic genes**

| RefSeq Transcript ID | Gene Symbol | Length | 5'UTR-<br>RBM3 binding site | CDS-<br>RBM3 binding site | 3'UTR-<br>RBM3 binding site |
|---|---|---|---|---|---|
| NM_009463 | Ucp1 | 5'UTR 1-231<br>CDS 232-1155<br>3'UTR 1156-1636 | GCAGGAG (118-124) | GGAGGTG (601-607) | none |
| NM_001308352 | PPARγ | 5'UTR 1-493<br>CDS 494-1921<br>3'UTR 1922-2125 | GCGGCAG (95-101) | GCGGCAG (1300-1306)<br>GGAGCAG (1726-1732) | none |
| NR_027710 | PGC1-α | 5'UTR 1-140<br>CDS 141-2534<br>3'UTR 2535-6464 | GGGGCTG (56-62)<br>GGAGCTG (133-139) | GCAGCAG (829-835)<br>GGAGCTG (1517-1523)<br>GCGGGAG (2105-2111) | GCTGCTG (2750-2756)<br>GGAGCAG (3526-3532)<br>GCTGGAG (4554-4560)<br>GCAGCAG (5970-5976) |
| NR_027710 | Prdm16 | 5'UTR 1-107<br>CDS 108-3938<br>3'UTR 3939-8601 | GGAGGAG (76-82)<br>GGAGGAG (79-85) | GGAGGAG (212-218)<br>GGAGCTG (384-390)<br>GGAGCAG (491-497)<br>GCAGCTG (604-610)<br>GCAGGAG (873-879)<br>GGGGGTG (2058-2064)<br>GCTGCTG (2153-2159)<br>GGAGCTG (2169-2175)<br>GCAGGAG (2243-2249)<br>GCAGCAG (2533-2539)<br>GGAGGTG (2559-2565)<br>GGGGCTG (2621-2617)<br>GCTGGAG (2831-2837)<br>GGAGCAG (3035-3041)<br>GCAGCAG (3164-3170)<br>GGAGGAG (3467-3473)<br>GGAGGAG (3479-3485)<br>GGAGCTG (3482-3488)<br>GCTGGAG (3485-3491)<br>GGAGGTG (3625-3631)<br>GGGGCTG (3689-3695)<br>GCAGCTG (3708-3714) | GCTGCAG (3982-3988)<br>GCAGGAG (3985-3991)<br>GGAGCAG (4008-4014)<br>GGAGCTG (4168-4174)<br>GCAGGAG (4230-4236)<br>GCTGCAG (4710-4716)<br>GCAGCAG (4875-4881)<br>GCAGCAG (6021-6027)<br>GCAGCAG (6348-6354)<br>GGTGGAG (7408-7414)<br>GGGGCTG (8085-8091)<br>GCTGGTG (8088-8094) |

### Supplementary Table 2: Statistical analysis for the qPCR experiments

**BAT qPCR Ucp1 (Fig. 3L)**

| Tukey's multiple comparisons test | Mean Diff. | 95.00% CI of diff. | Significant? | Summary | Adjusted P Value | |
|---|---|---|---|---|---|---|
| WT R.T. vs. KO R.T. | 0.9251 | 0.2980 to 1.552 | Yes | ** | 0.0027 | A-B |
| WT R.T. vs. WT cold | -1.239 | -1.866 to -0.6117 | Yes | *** | 0.0001 | A-C |
| WT R.T. vs. KO cold | 0.967 | 0.3400 to 1.594 | Yes | ** | 0.0018 | A-D |
| KO R.T. vs. WT cold | -2.164 | -2.791 to -1.537 | Yes | **** | <0.0001 | B-C |
| KO R.T. vs. KO cold | 0.04193 | -0.5851 to 0.6690 | No | ns | 0.9976 | B-D |
| WT cold vs. KO cold | 2.206 | 1.579 to 2.833 | Yes | **** | <0.0001 | C-D |

**BAT qPCR Pparγ (Fig. 3M)**

| Tukey's multiple comparisons test | Mean Diff. | 95.00% CI of diff. | Significant? | Summary | Adjusted P Value | |
|---|---|---|---|---|---|---|
| WT R.T. vs. KO R.T. | 0.9451 | 0.5028 to 1.387 | Yes | **** | <0.0001 | A-B |
| WT R.T. vs. WT cold | -0.7645 | -1.207 to -0.3222 | Yes | *** | 0.0005 | A-C |
| WT R.T. vs. KO cold | 0.9604 | 0.5181 to 1.403 | Yes | **** | <0.0001 | A-D |
| KO R.T. vs. WT cold | -1.71 | -2.152 to -1.267 | Yes | **** | <0.0001 | B-C |
| KO R.T. vs. KO cold | 0.01531 | -0.4270 to 0.4576 | No | ns | 0.9997 | B-D |
| WT cold vs. KO cold | 1.725 | 1.283 to 2.167 | Yes | **** | <0.0001 | C-D |

**BAT qPCR Pgc1α (Fig. 3N)**

| Tukey's multiple comparisons test | Mean Diff. | 95.00% CI of diff. | Significant? | Summary | Adjusted P Value | |
|---|---|---|---|---|---|---|
| WT R.T. vs. KO R.T. | 0.8648 | -4.020 to 5.749 | No | ns | 0.9592 | A-B |
| WT R.T. vs. WT cold | -22.6 | -27.49 to -17.72 | Yes | **** | <0.0001 | A-C |
| WT R.T. vs. KO cold | 0.3897 | -4.495 to 5.274 | No | ns | 0.9959 | A-D |
| KO R.T. vs. WT cold | -23.47 | -28.35 to -18.58 | Yes | **** | <0.0001 | B-C |
| KO R.T. vs. KO cold | -0.475 | -5.359 to 4.409 | No | ns | 0.9927 | B-D |
| WT cold vs. KO cold | 22.99 | 18.11 to 27.88 | Yes | **** | <0.0001 | C-D |

**BAT qPCR Prdm16 (Fig. 3O)**

| Tukey's multiple comparisons test | Mean Diff. | 95.00% CI of diff. | Significant? | Summary | Adjusted P Value | |
|---|---|---|---|---|---|---|
| WT R.T. vs. KO R.T. | 0.6074 | -1.003 to 2.218 | No | ns | 0.6003 | A-B |
| WT R.T. vs. WT cold | -2.232 | -3.842 to -0.6211 | Yes | * | 0.007 | A-C |
| KO R.T. vs. WT cold | -2.839 | -4.449 to -1.229 | Yes | ** | 0.001 | B-C |

**RIP qPCR Ucp1 (Fig. 4F)**

| Tukey's multiple comparisons test | Mean Diff. | 95.00% CI of diff. | Significant? | Summary | Adjusted P Value | |
|---|---|---|---|---|---|---|
| WT IgG IP vs. WT RBM3 IP | -2.881 | -5.381 to -0.3810 | Yes | * | 0.0203 | A-B |
| WT IgG IP vs. KO IgG IP | 1.67E-09 | -2.500 to 2.500 | No | ns | >0.9999 | A-C |
| WT IgG IP vs. KO RBM3 IP | -0.5807 | -3.081 to 1.920 | No | ns | 0.9143 | A-D |
| WT RBM3 IP vs. KO IgG IP | 2.881 | 0.3810 to 5.381 | Yes | * | 0.0203 | B-C |
| WT RBM3 IP vs. KO RBM3 IP | 2.3 | -0.1998 to 4.801 | No | ns | 0.0782 | B-D |
| KO IgG IP vs. KO RBM3 IP | -0.5807 | -3.081 to 1.920 | No | ns | 0.9143 | C-D |

**RIP qPCR Pparγ (Fig. 4G)**

| Tukey's multiple comparisons test | Mean Diff. | 95.00% CI of diff. | Significant? | Summary | Adjusted P Value | |
|---|---|---|---|---|---|---|
| WT IgG IP vs. WT RBM3 IP | -5.464 | -13.94 to 3.014 | No | ns | 0.2673 | A-B |
| WT IgG IP vs. KO IgG IP | 2.00E-09 | -15.34 to 15.34 | No | ns | >0.9999 | A-C |
| WT IgG IP vs. KO RBM3 IP | -0.2012 | -10.43 to 10.02 | No | ns | >0.9999 | A-D |
| WT RBM3 IP vs. KO IgG IP | 5.464 | -9.659 to 20.59 | No | ns | 0.704 | B-C |
| WT RBM3 IP vs. KO RBM3 IP | 5.263 | -4.637 to 15.16 | No | ns | 0.4175 | B-D |
| KO IgG IP vs. KO RBM3 IP | -0.2012 | -16.37 to 15.97 | No | ns | >0.9999 | C-D |

**RIP qPCR Pgc1α (Fig. 4H)**

| Tukey's multiple comparisons test | Mean Diff. | 95.00% CI of diff. | Significant? | Summary | Adjusted P Value | |
|---|---|---|---|---|---|---|
| WT IgG IP vs. WT RBM3 IP | -1.09 | -3.814 to 1.635 | No | ns | 0.6761 | A-B |
| WT IgG IP vs. KO IgG IP | 0 | -2.724 to 2.724 | No | ns | >0.9999 | A-C |
| WT IgG IP vs. KO RBM3 IP | 0.8861 | -1.959 to 3.732 | No | ns | 0.8151 | A-D |
| WT RBM3 IP vs. KO IgG IP | 1.09 | -1.508 to 3.687 | No | ns | 0.6433 | B-C |
| WT RBM3 IP vs. KO RBM3 IP | 1.976 | -0.7486 to 4.700 | No | ns | 0.2072 | B-D |
| KO IgG IP vs. KO RBM3 IP | 0.8861 | -1.838 to 3.611 | No | ns | 0.795 | C-D |

**RIP qPCR Prdm16 (Fig. 4I)**

| Tukey's multiple comparisons test | Mean Diff. | 95.00% CI of diff. | Significant? | Summary | Adjusted P Value | |
|---|---|---|---|---|---|---|
| WT IgG IP vs. WT RBM3 IP | -0.8442 | -3.403 to 1.715 | No | ns | 0.7635 | A-B |
| WT IgG IP vs. KO IgG IP | 0 | -2.369 to 2.369 | No | ns | >0.9999 | A-C |
| WT IgG IP vs. KO RBM3 IP | -0.1631 | -2.899 to 2.573 | No | ns | 0.9979 | A-D |
| WT RBM3 IP vs. KO IgG IP | 0.8442 | -1.319 to 3.007 | No | ns | 0.6623 | B-C |
| WT RBM3 IP vs. KO RBM3 IP | 0.6811 | -1.878 to 3.240 | No | ns | 0.8575 | B-D |
| KO IgG IP vs. KO RBM3 IP | -0.1631 | -2.532 to 2.206 | No | ns | 0.9968 | C-D |
